# Molecular crowding differentially regulates magnetic field-dependent fluorescence, recovery kinetics, and photostability of the magnetosensitive protein MagLOV2

**DOI:** 10.64898/2026.10.30.762980

**Authors:** Venkatesh Sridharan, Brian L. Ross, Alessandro Lodesani

## Abstract

Macromolecular crowding and solvent composition can alter protein conformation, diffusion, hydration, and cofactor photophysics, yet their influence on magnetically sensitive flavoproteins is poorly understood. We tested seven molecular environments in MagLOV2-overexpressing *Escherichia coli* lysate using a custom-built, high-intensity magnetofluorescence platform. Five-point concentration series of Dextran 70, Ficoll PM70, PEG6000, PEG3350, glycerol, ethylene glycol, and bovine serum albumin (BSA) were measured at 22 mT under continuous 6.12 W/cm^2^ blue-light excitation. Dextran 70, Ficoll PM70, and PEG6000 progressively attenuated the negative magnetic field effect (MFE), whereas PEG3350 shifted toward mild enhancement at higher concentrations. Glycerol and ethylene glycol enhanced the negative MFE despite producing the strongest slowing of recovery upon the field being switched off (“OFF-recovery”), while BSA modestly enhanced MFE and accelerated OFF-recovery. Across 105 treatment cuvettes, the change in MFE was not globally correlated with changes in the OFF-recovery time constant (*r* = −0.07, *p* = 0.449). The initial pre-field illumination period revealed a second phenotype: Dextran and Ficoll increased late fluorescence loss, whereas glycerol and BSA were photoprotective. Pre-field fluorescence loss correlated moderately with MFE change across the full dataset (*r* = 0.474, *p* = 3.28 × 10^−7^) but not consistently within crowder classes. These findings show that MagLOV2 spin-dependent magnetofluorescence is shaped by crowder chemistry, polymer size, solvent environment, recovery kinetics, and photostability rather than by a single generic crowding or viscosity mechanism.

## 1. Introduction

Proteins are commonly characterized in dilute aqueous buffers, whereas cellular proteins function in a highly volume-occupied and chemically heterogeneous environment. Proteins, nucleic acids, ribosomes, carbohydrates, and other macromolecules can occupy approximately 10–40% of cellular volume, restricting the space accessible to individual biomolecules [1, 2]. A central physical consequence is excluded volume: extended conformations occupy more effective volume than compact conformations, so crowding can favor protein compaction and thermodynamic stability. In cells and concentrated model solutions, however, steric effects coexist with weak chemical interactions, altered hydration, and changes in translational and rotational diffusion [1–3].

Crowding therefore does not uniformly increase protein function. Stabilization may improve activity when a rigid native state is favorable, but it can reduce activity when catalysis or signaling requires conformational exchange, domain motion, substrate transport, or cofactor access. Intrinsically disordered regions are particularly sensitive because their broad conformational ensembles can compact, remain relatively unchanged, or expand depending on sequence and crowder chemistry [4]. Crowding also modifies intermolecular association, aggregation, and liquid–liquid phase separation, linking the physical intracellular environment to biomolecular condensates and disease-associated protein assembly [5]. Sickle hemoglobin polymerization provides a clear example in which very high intracellular protein concentration and nonideal-solution behavior amplify a disease-causing self-association process [6]. Finally, molecular crowding can substantially alter spin chemistry by changing molecular diffusion, radical-pair separation, recombination kinetics, and the local environment of unpaired electron spins.

Agents used to emulate crowding are themselves mechanistically distinct. Dextran and Ficoll mainly probe polymeric excluded volume but differ in flexibility and architecture; PEG combines excluded-volume effects with molecular weight-dependent soft interactions; glycerol and ethylene glycol are better considered small-molecule cosolvents or microviscogens that alter viscosity, hydration, hydrogen bonding, and osmotic properties; and BSA provides a proteinaceous crowded environment with heterogeneous interaction surfaces [3, 7–10]. Consequently, the identity and size of a crowding agent can be as important as its concentration, and dilute biochemical assays may not fully predict protein behavior in cellular-like environments.

Whether these environmental effects influence protein magnetosensitivity remains largely unexplored. To our knowledge, no published study has systematically examined how macromolecular crowding modifies the underlying spin physics, and consequently the magnetic field-dependent fluorescence, of a magnetosensitive protein. MagLOV2 is an engineered LOV-domain flavoprotein derived by directed evolution from the C450A variant of the *Avena sativa* phototropin 1 LOV2 domain. MagLOV2 exhibits a large fluorescence magnetic field effect (MFE) both when purified and in living bacterial cells. Magnetic resonance measurements support a spin-correlated radical-pair mechanism involving the flavin cofactor and its protein environment [11]. Independent field-strength measurements further revealed a non-monotonic response with a low-field sign reversal, consistent with radical-pair spin chemistry [12].

The structural scaffold provides a plausible route by which the molecular environment could tune this response. AsLOV2 contains an FMN-binding PAS domain core coupled to the N-terminal A^′^*α* and C-terminal J*α* helices. Structural and time-resolved spectroscopic studies show that perturbations originating around FMN propagate through the *β*-sheet scaffold and drive changes in the A^′^*α*/J*α* regions [13–15]. The three-dimensional structure and conformational landscape of MagLOV2 itself have not yet been experimentally resolved, so extrapolation from AsLOV2 should remain mechanistic rather than structural proof. Nevertheless, crowding could alter protein compaction, helix docking, local hydration, dielectric properties, oxygen accessibility, and the geometry or dynamics of residues surrounding FMN.

These environmental effects may influence more than MFE amplitude. A recent four-state kinetic model of fluorescent flavoprotein magnetosensitivity predicts that the observed fluorescence contrast depends on the balance between photoexcitation and slower spin-independent recovery reactions, not solely on radical-pair singlet yield [16]. Crowder-dependent changes in protein dynamics, solvent organization, or flavin photochemistry could therefore alter kinetics upon the field being switched on (“ON-kinetics”), OFF-recovery, photostability, and the apparent steady-state MFE through partially separable mechanisms.

Here, we tested this hypothesis using MagLOV2-overexpressing *Escherichia coli* (*E. coli*) lysate exposed to five concentrations of seven mechanistically distinct molecular environments: Dextran 70, Ficoll PM70, PEG3350, PEG6000, glycerol, ethylene glycol, and BSA. We quantified steady-state MFE amplitude, field-ON and field-OFF kinetics, reversibility, and pre-field illumination-dependent fluorescence loss. The design was intended to determine whether MagLOV2 magnetofluorescence follows a generic crowding/viscosity axis or instead reports crowder-specific effects on protein dynamics, solvent interactions, and the FMN microenvironment.

## 2. Methods

### 2.1 MagLOV2 expression and lysate preparation

*E. coli* BL21(DE3) carrying pRSET-MagLOV2 was used for recombinant MagLOV2 production. Expression was induced with 100 *µ*M isopropyl *β*-D-1-thiogalactopyranoside (IPTG) for 6 h at 37 °C in the presence of 500 *µ*M riboflavin. Cell pellets were lysed using freshly prepared 1× BugBuster Protein Extraction Reagent supplemented with lysozyme and Benzonase. For a pellet from approximately 100 mL culture, 2–5 mL lysis mixture was used depending on pellet size. Lysozyme was used at approximately 0.5–1.0 mg/mL, and Benzonase was added according to the supplier recommendation (approximately 25–30 U/mL in the concentrated lysis suspension as a working range). EDTA-free protease inhibitors were used when required. MagLOV2 was evaluated as a crude bacterial lysate rather than as purified protein. The same lysate batch was used throughout the confirmatory study to minimize variation in MagLOV2 abundance and lysate composition.

### 2.2 Sample preparation and molecular environments

Each cuvette contained 100 *µ*L MagLOV2-overexpressing *E. coli* lysate and 700 *µ*L buffer/crowder solution, corresponding to an eightfold final lysate dilution. Samples were prepared in 50 mM Tris-HCl, pH 7.4. Dextran 70 and Ficoll PM70 were tested at 25, 50, 100, 150, and 200 mg/mL; PEG6000 and PEG3350 at 12.5, 25, 50, 75, and 100 mg/mL; glycerol and ethylene glycol at 2.5, 5, 10, 15, and 20% (v/v); and BSA at 10, 15, 20, 25, and 30 mg/mL. The two PEGs are chemically identical and differ only in chain length (average molecular weights of approximately 6000 and 3350 g/mol), so their comparison isolates the effect of polymer size. For each condition, three fresh cuvettes were prepared independently and measured sequentially. Repeated field cycles within a cuvette were technical measurements, not independent replicates. Pilot experiments were used only to establish acquisition and analysis parameters and were not pooled with the five-point confirmatory dataset.

### 2.3 High-intensity magnetofluorescence instrumentation

Magnetofluorescence measurements were performed with a custom, high-intensity, cuvette-based magnetofluorescence screener developed in-house. Samples were positioned in an Ocean Optics SQ1-ALL cuvette holder within the uniform field region of a water-cooled Helmholtz-coil assembly. Blue excitation (465 nm laser) was delivered through a multimode optical-fiber pathway, with laser output controlled by analog voltage from a National Instruments data-acquisition (DAQ) system. Illumination intensity at the sample was calibrated from optical-power and beam-profile measurements; the platform was capable of intensities up to approximately 25.8 W/cm^2^.

Fluorescence was detected in free space using a Thorlabs FDS100 silicon photodiode coupled to a Thorlabs PDA200C amplifier. Magnetic fields were generated by four subcoils configured in a 2-series/2-parallel arrangement and driven by a GW Instek ASR-2100 programmable AC/DC source under DAQ analog control. The measured field was recorded continuously with a Texas Instruments DRV5055A2 analog Hall-effect sensor aligned to the sample field. Laser control, field switching, fluorescence acquisition, and sensor acquisition were synchronized through custom Python-based software.

### 2.4 Magnetofluorescence acquisition

Measurements were performed at 25 °C under continuous blue-light (465 nm) excitation at 6.12 W/cm^2^. The target magnetic field was 22 mT, and the simultaneously measured Hall-sensor trace was retained for each run. Each acquisition began with a 10 s field-OFF (B-OFF) interval, followed by five cycles of 10 s field ON and 15 s field OFF. For kinetic fitting, the moments the field switched on and off were taken from the measured Hall-sensor trace rather than from the times at which the software issued the switching commands.

### 2.5 Primary MFE and kinetic analysis

Instrument-conditioned fluorescence and field traces were analyzed after despiking/rolling averaging and reduction to approximately 100 Hz. Because fluorescence declined during continuous illumination, a time-dependent zero-field baseline was estimated using piecewise cubic Hermite interpolating polynomial (PCHIP) interpolation through 1 s median anchors from late B-OFF intervals after recovery. A global exponential photobleaching fit to the same OFF anchors was retained as a sensitivity analysis.

For cycle *c*, baseline-normalized residual fluorescence was calculated as

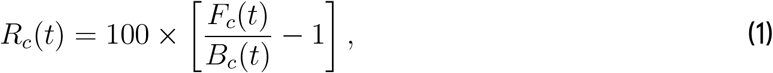

where *F*_*c*_(*t*) is the measured fluorescence and *B*_*c*_(*t*) is the estimated zero-field baseline. The magnetic field effect was equivalently expressed as

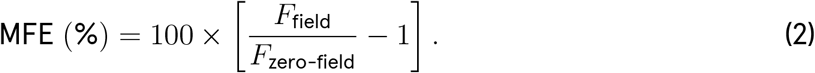

Negative values therefore indicate a field-associated decrease in fluorescence. At a field of 22 mT, this is most likely consistent with a triplet-born radical pair. The primary steady-state endpoint was the residual averaged over the window from 6 to 9 s after the sensor-defined field-ON transition. Cycle 1 was retained for adaptation and quality control; the cuvette-level primary endpoint was the mean of cycles 2–5:

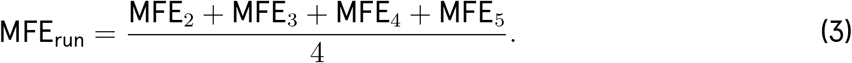

Field-ON and field-OFF transitions were fitted empirically with single-exponential models. ON kinetics were described by

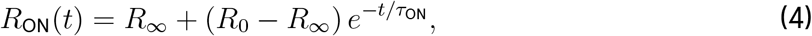

and OFF recovery by

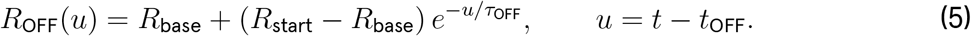

The corresponding half-times were *t*_1/2,ON_ = *τ*_ON_ ln 2 and *t*_1/2,OFF_ = *τ*_OFF_ ln 2. Percentage recovery after an OFF interval *T* was calculated as

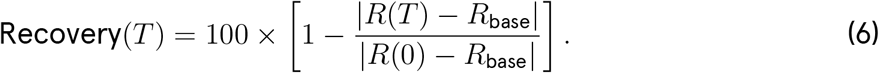

The fitted *τ*_ON_ and *τ*_OFF_ values were treated as seconds-scale empirical fluorescence/protein-state kinetics and not as direct measurements of radical-pair spin lifetimes.

### 2.6 Pre-field fluorescence-loss analysis

The immediate laser-on peak reached the detector ceiling in 95 of 111 lysate-bearing runs and was therefore not used for quantitative between-condition comparison. The primary pre-field endpoint captured the later nonsaturated B-OFF phase: *F*_ref_ was the mean fluorescence at 6.0–6.5 s, and *F*_pre-field_ was the mean from 1.0 to 0.25 s before the first field pulse. Late-phase fluorescence loss was calculated as 100 × (*F*_ref_ − *F*_pre-field_)/*F*_ref_. This endpoint is referred to operationally as pre-field photobleaching, while recognizing that LOV fluorescence loss can also include reversible flavin photoreduction or dark-state accumulation [17].

### 2.7 Local-control adjustment, statistics, and quality control

Each independently prepared cuvette was the experimental unit (*n* = 3 per concentration). Buffer-only control cuvettes were measured periodically throughout the acquisition series. Each treatment cuvette was compared with a local control: the control MFE and *τ*_OFF_ expected at that point in the series, interpolated from the nearest complete controls measured before and after it. ΔMFE was defined as treatment MFE minus local-control MFE, expressed in percentage points; because the underlying MFE is negative, positive ΔMFE indicates attenuation and negative ΔMFE indicates enhancement. Recovery changes were expressed as

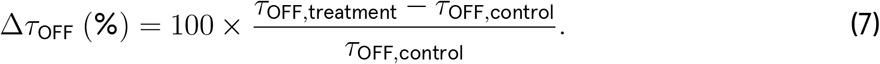

Agent-specific concentration models included concentration and replicate, and matched-agent comparisons used agent × concentration interaction models. Holm correction [18] was applied across agent-specific concentration-trend tests. Pearson correlations were calculated on cuvette-level values. Effect size and replicate consistency were emphasized because all cuvettes originated from a single lysate batch. No point-by-point blank subtraction was applied; no-lysate blanks were retained as optical and field-coupled control measurements.

## 3. Results

### 3.1 Crowder identity and concentration reshape the MagLOV2 magnetic response

Buffer-control MFE ranged from −54.84% to −51.83% across the acquisition series. Field delivery and within-cuvette repeatability were strong: all treatment runs remained within ±5% of the 22 mT target, the median cycle coefficient of variation (CV) was 0.68%, and only 1 of 105 treatment cuvettes exceeded 5% CV. Against this stable assay background, molecular-environment effects were strongly agent dependent (Table I).

**Table 1.** Highest-concentration local control-adjusted MagLOV2 response phenotypes. ΔMFE is in percentage points relative to the local control.

| Agent | Highest dose | ΔMFE (pp) | Δτ<sub>OFF</sub> (%) | Phenotype |
|---|---|---|---|---|
| Dextran 70 | 200 mg/mL | +7.65 ± 0.34 | +11.2 ± 1.1 | Strong attenuation; slower recovery |
| Ficoll PM70 | 200 mg/mL | +4.05 ± 1.28 | +15.0 ± 3.6 | Moderate attenuation; slower recovery |
| PEG6000 | 100 mg/mL | +6.77 ± 0.05 | +8.9 ± 0.8 | Strong attenuation; slower recovery |
| PEG3350 | 100 mg/mL | −0.95 ± 0.22 | +6.0 ± 0.7 | Mild enhancement; slower recovery |
| Glycerol | 20% v/v | −1.84 ± 1.14 | +20.3 ± 2.7 | Enhancement; strongly slower recovery |
| Ethylene glycol | 20% v/v | −2.55 ± 0.48 | +24.6 ± 1.2 | Enhancement; strongest slowing |
| BSA | 30 mg/mL | −1.10 ± 0.15 | −8.0 ± 1.6 | Mild enhancement; faster recovery |

Dextran 70, Ficoll PM70, and PEG6000 progressively attenuated the negative MFE (Fig. 1). At the highest concentrations, Dextran 70 and PEG6000 produced the largest shifts (+7.65 and +6.77 percentage points, respectively), whereas Ficoll PM70 produced a smaller attenuation (+4.05 percentage points). Dextran showed the predicted stronger concentration dependence than Ficoll, although the agent × concentration interaction was borderline (*p* = 0.058). PEG molecular weight was a major discriminator: PEG6000 attenuated MFE, whereas PEG3350 crossed toward mild enhancement at ≥ 50 mg/mL, with a strong PEG-size interaction (*p* = 8.95 × 10^−7^). Glycerol and ethylene glycol generally shifted the response toward a more negative MFE at higher concentrations, while BSA produced a smaller enhancement that plateaued at 20–30 mg/mL.

**Fig. 1.**
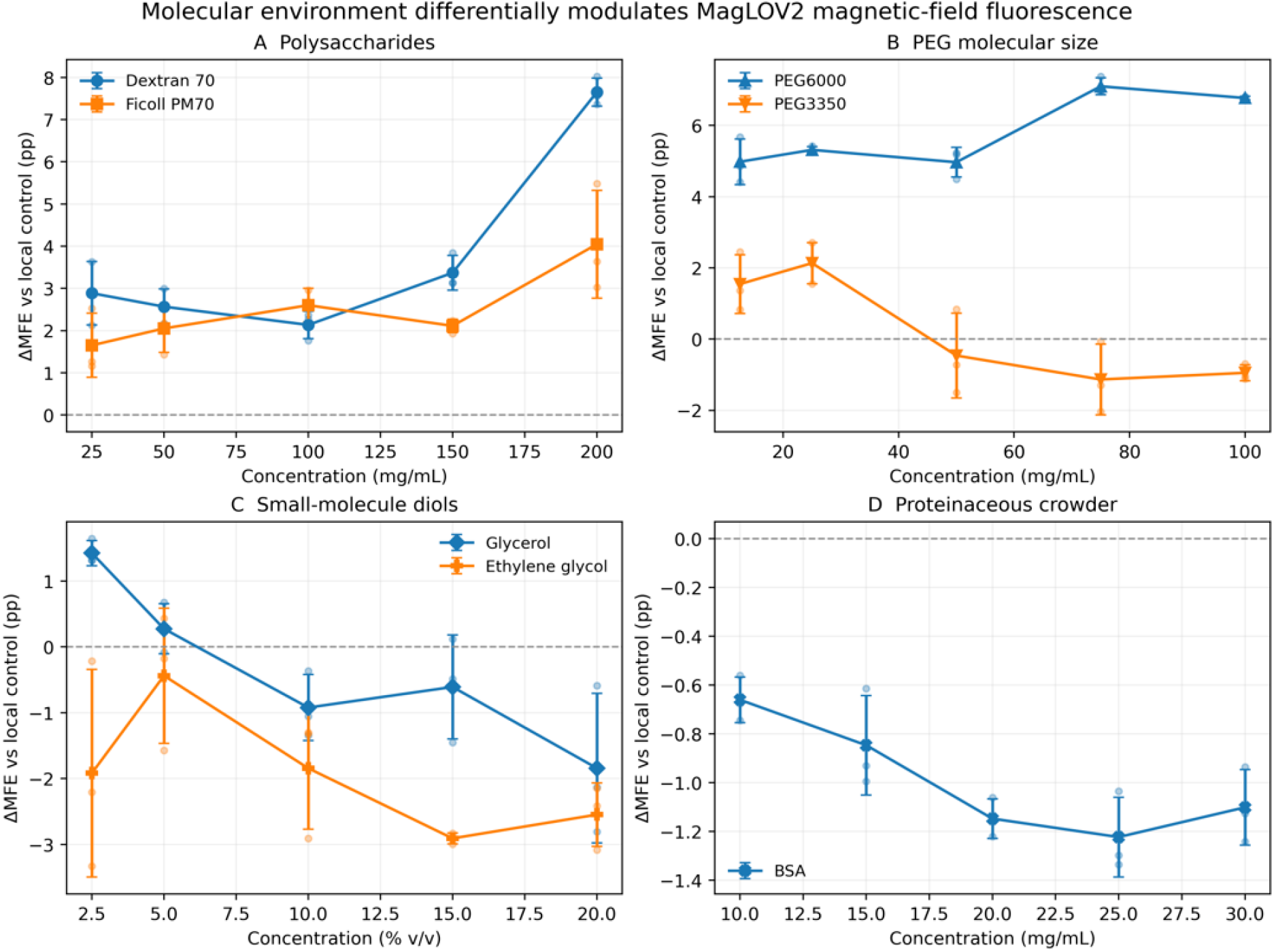
Local control-adjusted steady-state magnetic field effect across the five-point concentration series. A Polysaccharide crowding agents (Dextran 70 and Ficoll PM70); B polyethylene glycol (PEG), both 3350 and 6000 molecular weight; C small-molecule diols (glycerol and ethylene glycol); and D bovine serum albumin (BSA), a protein crowder. Positive ΔMFE indicates attenuation of the negative field-associated fluorescence change; negative ΔMFE indicates enhancement. Points represent the average values plotted with their standard deviations; faint points are individual fresh cuvettes.

### 3.2 OFF-recovery kinetics are strongly altered but do not globally determine MFE amplitude

Dextran, Ficoll, both PEGs, glycerol, and ethylene glycol slowed field-OFF recovery with increasing concentration (Fig. 2). The largest changes occurred with the small-molecule diols: *τ*_OFF_ increased by approximately 20% with 20% glycerol and 25% with 20% ethylene glycol. BSA moved in the opposite direction, accelerating recovery by approximately 8% at 30 mg/mL. Despite this broad kinetic sensitivity, ΔMFE and Δ*τ*_OFF_ were essentially uncorrelated across all 105 treatment cuvettes (Pearson *r* = −0.07, *p* = 0.449). Coupling was agent-class dependent: Dextran, Ficoll, and PEG6000 generally showed slower recovery together with MFE attenuation, whereas glycerol and ethylene glycol showed slower recovery together with MFE enhancement. Thus, spin-independent recovery is environmentally sensitive but cannot by itself explain MFE amplitude.

**Fig. 2.**
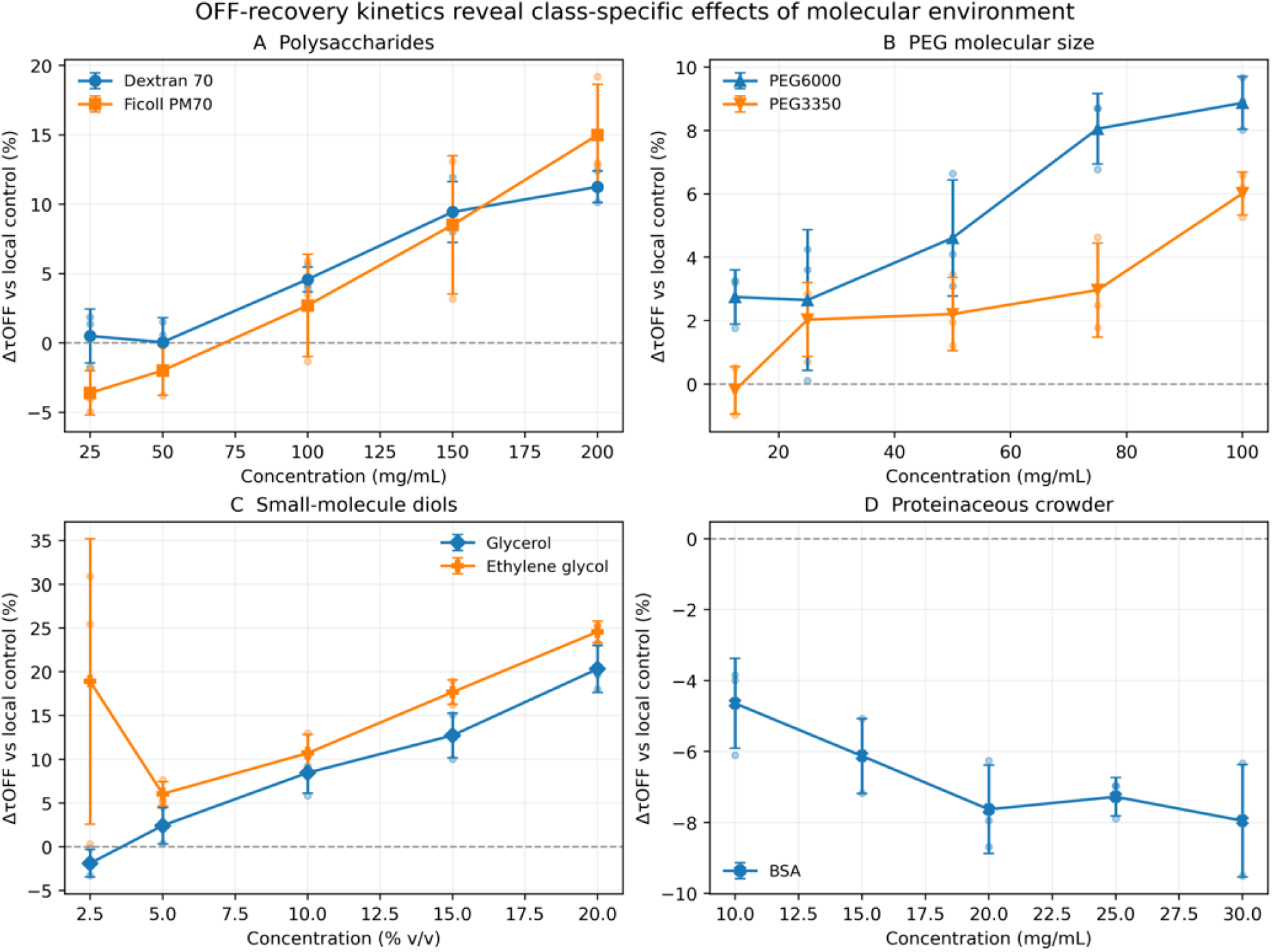
Local control-adjusted field-OFF recovery time constant (*τ*_OFF_) for MagLOV2 across the five-point concentration series. A Polysaccharide crowding agents (Dextran 70 and Ficoll PM70); B polyethylene glycol (PEG), both 3350 and 6000 molecular weight; C small-molecule diols (glycerol and ethylene glycol); and D bovine serum albumin (BSA), a protein crowder. Positive values indicate slower recovery and negative values faster recovery. Points represent the average values plotted with their standard deviations; faint points are individual cuvettes.

### 3.3 The initial B-OFF interval reveals a distinct photostability phenotype

The molecular environment also altered fluorescence persistence before magnetic-field exposure (Fig. 3). At the highest concentrations, Dextran 70 (+2.93 ± 0.38 percentage points) and Ficoll PM70 (+3.34 ± 0.66 percentage points) increased late fluorescence loss, whereas glycerol (−2.25 *±* 0.33 percentage points) and BSA (−2.27 ± 0.35 percentage points) were strongly photoprotective. PEG3350 showed a concentration-dependent crossover from protection to mild pro-bleaching (+0.91 ± 0.25 percentage points at 100 mg/mL), PEG6000 was approximately neutral at 100 mg/mL (−0.14 ± 0.33 percentage points), and ethylene glycol showed mild protection (−0.47 ± 0.47 percentage points at 20%).

**Fig. 3.**
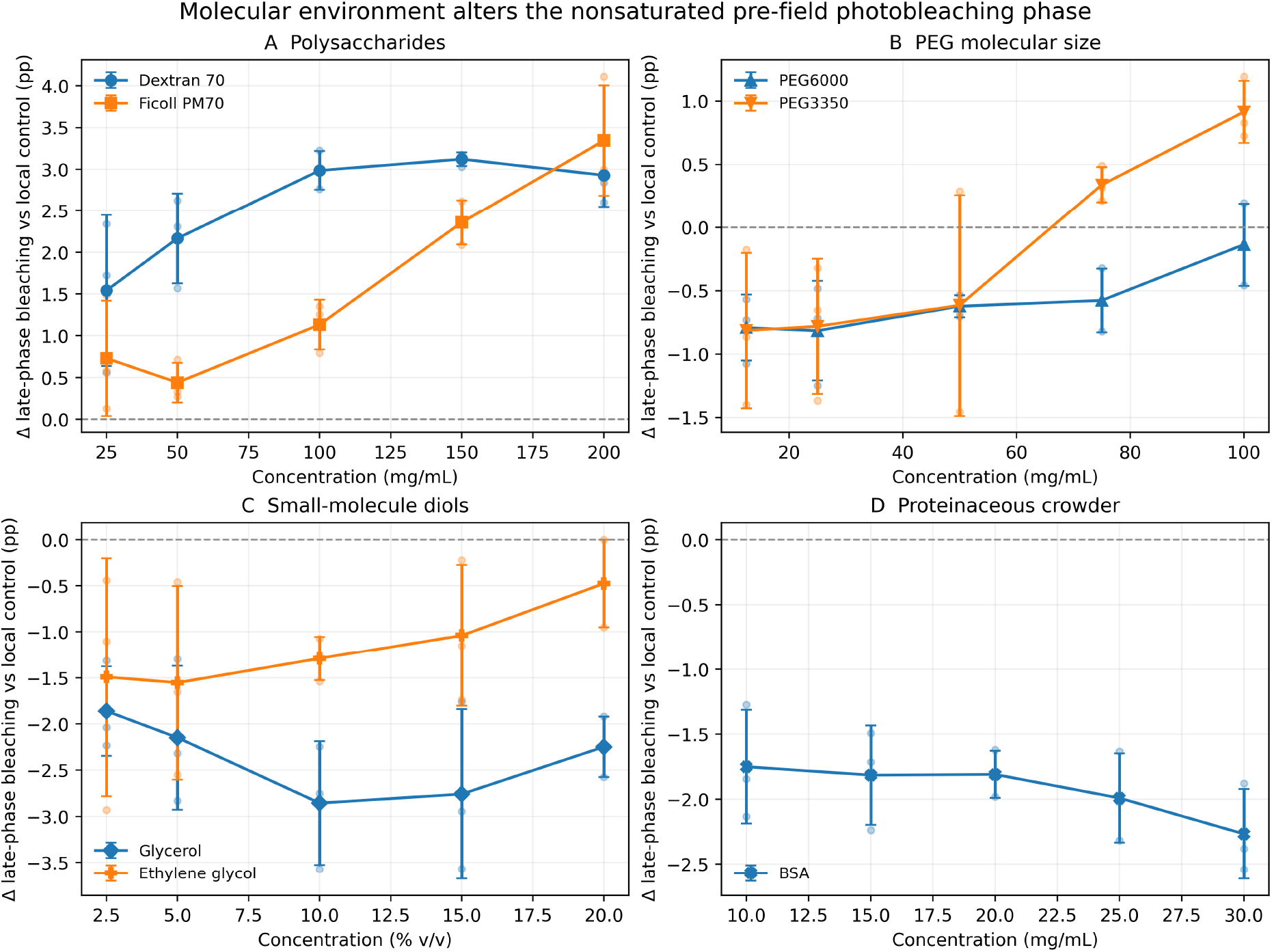
Local control-adjusted late pre-field fluorescence loss during the initial B-OFF illumination period. A Polysaccharide crowding agents (Dextran 70 and Ficoll PM70); B polyethylene glycol (PEG), both 3350 and 6000 molecular weight; C small-molecule diols (glycerol and ethylene glycol); and D bovine serum albumin (BSA), a protein crowder. Positive values indicate greater fluorescence loss; negative values indicate photoprotection. Points represent the average values plotted with their standard deviations; faint points are individual cuvettes.

Across the full dataset, greater pre-field fluorescence loss was moderately associated with attenuation of the negative MFE (Fig. 4; Pearson *r* = 0.474, *p* = 3.28 × 10^−7^). This relationship was not uniform by agent: Ficoll showed positive coupling, whereas PEG3350 and ethylene glycol showed strong negative within-agent relationships. Pre-field fluorescence loss did not significantly correlate with Δ*τ*_OFF_ (*r* = 0.142, *p* = 0.149). MFE amplitude, OFF-recovery, and photostability therefore represent partially separable outputs of the same environment-sensitive flavoprotein system.

**Fig. 4.**
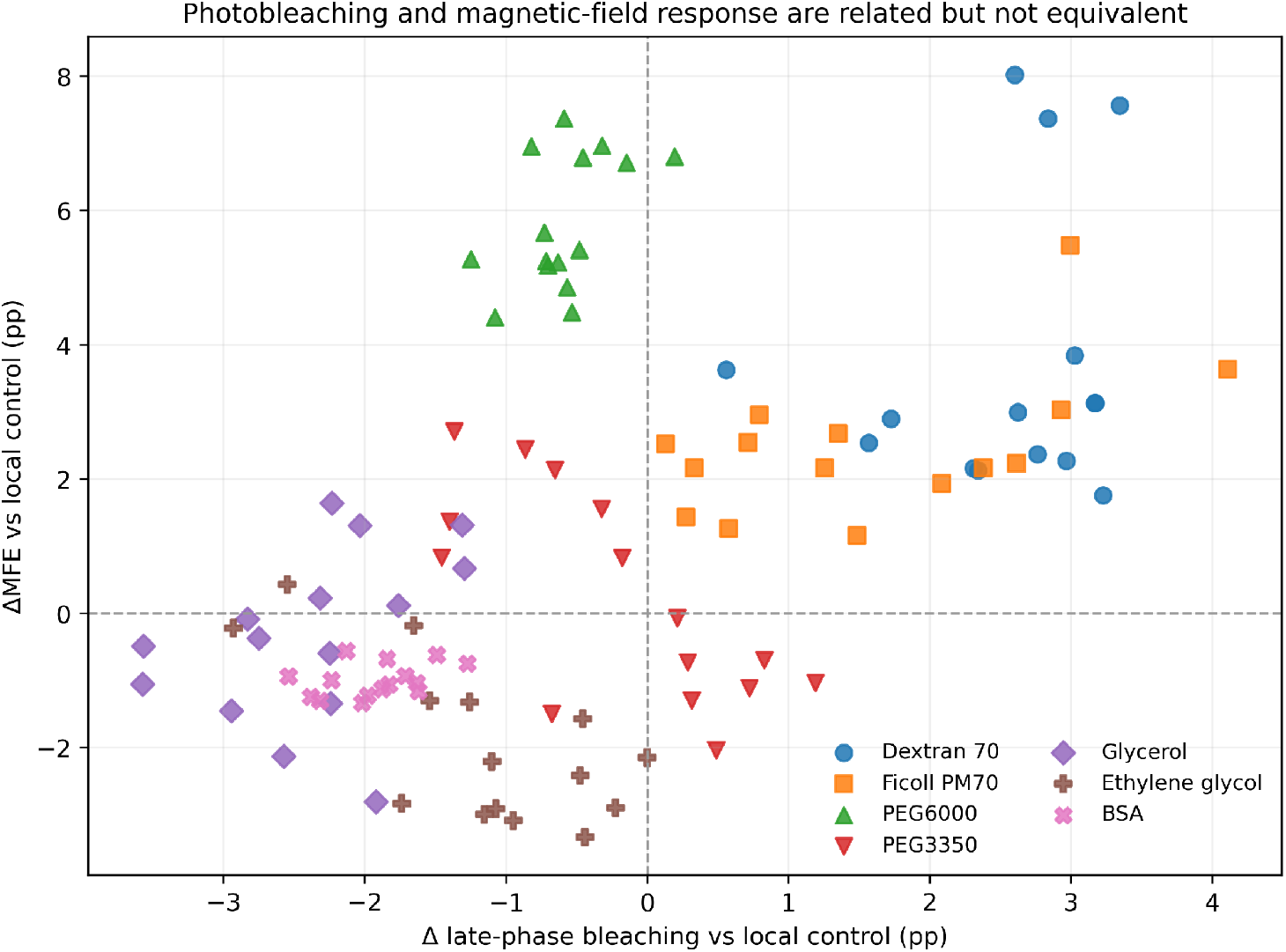
Relationship between local control-adjusted late pre-field fluorescence loss and local control-adjusted MFE across individual treatment cuvettes.

## 4. Discussion

This study demonstrates that the molecular environment is a major regulator of MagLOV2 magnetic field-dependent fluorescence, recovery kinetics, and photostability, but the response cannot be reduced to a single generic measure of crowding. Dextran 70, Ficoll PM70, and PEG6000 attenuated the negative MFE, whereas the small-molecule diols generally enhanced it at higher concentration. PEG3350 diverged sharply from PEG6000 despite identical chemical repeat units, and BSA produced a distinct proteinaceous-crowding phenotype with mild MFE enhancement, faster recovery, and photoprotection. Together, these results establish crowder chemistry and molecular size as mechanistic variables rather than interchangeable ways to increase solution viscosity or occupied volume.

The findings are consistent with the modern crowding framework in which excluded volume competes with weak chemical interactions, hydration, and changes in conformational dynamics [1–3]. Dextran and Ficoll differ in architecture and flexibility, and carbohydrate crowders of different size and shape can stabilize proteins to different extents while altering function [7, 9]. PEG is particularly nonideal: its molecular weight can shift the balance between excluded-volume effects and weak protein–polymer interactions, producing qualitatively different conformational responses [8]. The opposing PEG3350 and PEG6000 MFE trends, together with their different pre-field fluorescence-loss profiles, therefore argue against using PEG concentration as a universal proxy for crowding strength.

The diol data provide a complementary separation between macromolecular crowding and small-solute effects. Glycerol and ethylene glycol produced the strongest slowing of OFF recovery while shifting MFE in the opposite direction from Dextran, Ficoll, and PEG6000. Low- and high-molecular-mass crowding agents are known to affect reaction kinetics differently [10], and the present results indicate that bulk viscosity alone is insufficient to explain MagLOV2 behavior. Solvent structure, preferential hydration, osmolarity, hydrogen-bonding networks, oxygen transport, and local dielectric properties are plausible additional variables. For MagLOV2, the relevant cofactor is noncovalently bound FMN, so local FMN mobility and solvent accessibility are likely more important than bulk cofactor diffusion; in other flavoproteins, diffusional access of redox partners or exchangeable cofactors could provide an additional environmental control.

These effects are mechanistically plausible within the LOV2 structural scaffold. The FMN-binding PAS core is coupled to conformationally dynamic A^′^*α* and J*α* regions, and light-induced perturbations around FMN propagate into helix rearrangement and unfolding [13–15]. Crowding could shift this conformational ensemble, alter the dynamics of the *β*-sheet/helix network, or reorganize water and hydrogen bonds around FMN. Even without a major change in global fold, small changes in side-chain orientation, hydration, oxygen accessibility, or donor–acceptor geometry could change electron-transfer branching, radical-pair formation and recombination, and recovery pathways. Because the structure of MagLOV2 itself has not yet been resolved, these possibilities should be treated as testable mechanisms rather than direct structural conclusions.

The pre-field fluorescence-loss phenotype adds a second photochemical dimension. Cysteine-less LOV proteins can undergo flavin electron transfer and photoreduction to semiquinone states [17]. Continuous high-intensity excitation may therefore redistribute FMN among fluorescent, dark, photoreduced, and irreversibly damaged states. The moderate global association between late fluorescence loss and MFE suggests that these outputs share upstream photochemistry, but the opposite within-agent relationships for PEG3350 and ethylene glycol demonstrate that fluorescence loss is not the sole determinant of magnetic sensitivity.

The kinetic results further refine interpretation of the radical-pair mechanism. A recently proposed four-state model predicts that fluorescence is a faithful reporter of magnetic field-dependent singlet yield only in kinetic regimes where photoexcitation competes effectively with slower spin-independent recovery [16]. Our data support the importance of this recovery axis because *τ*_OFF_ changes substantially with the molecular environment, yet the absence of a global MFE–*τ*_OFF_ relationship shows that recovery is not the only environmental control parameter. A more complete description of magnetofluorescence will likely require the coupled effects of photoexcitation, radical-pair branching, spin-independent recovery, conformational relaxation, and partitioning of FMN among fluorescent and nonfluorescent states.

These observations have practical implications for discovery and characterization of new magnetosensitive proteins. Magnetofluorescence assays are usually optimized around field strength, excitation intensity, detector sensitivity, and timing; the present study indicates that the physicochemical environment should be treated as an important assay dimension. A candidate protein may display a weak or undetectable MFE in one buffer yet a larger response under conditions that favor a productive conformation, cofactor microenvironment, or recovery regime. Conversely, the condition that maximizes basal fluorescence may not maximize magnetic sensitivity. A small, standardized matrix of polymeric crowders, cosolvents, osmolytes, and proteinaceous crowders could therefore help identify hidden or condition-dependent MFE phenotypes and define assay windows with improved dynamic range and kinetic resolution.

The same logic can be incorporated into directed evolution. Current selection strategies can enrich variants for MFE amplitude, response speed, fluorescence intensity, or photostability under a single environment [11]. Screening selected variants across controlled molecular environments would distinguish mutations that are robust to crowding from variants whose performance depends strongly on solvent coupling or conformational dynamics. Selection could also explicitly target high MFE under crowded conditions, rapid reversible kinetics, low photobleaching, or consistent responses across multiple environments. Simultaneous measurement of MFE amplitude, *τ*_ON_, *τ*_OFF_, and photostability thus provides a multidimensional phenotype for ranking engineered variants and for mapping which mutations primarily affect spin-sensitive chemistry versus the surrounding protein/solvent landscape.

More broadly, the results emphasize that radical-pair spin dynamics define the magnetic-field-sensitive step but do not alone determine the magnitude of the measured fluorescence phenotype. Protein conformational dynamics, local water organization, hydrogen bonding, osmolarity, oxygen transport, diffusional access of reaction partners, and cofactor microenvironment can alter the populations and kinetics of states that participate in or compete with the radical-pair pathway. Molecular crowding is therefore not merely a confounder in magnetobiology assays; it can serve as a controlled perturbation for identifying the physical variables that gate expression of magnetosensitivity.

Several limitations remain. All independent cuvette preparations originated from one MagLOV2 lysate batch, and crowders may influence endogenous lysate components in addition to MagLOV2. Concentration series were acquired sequentially rather than as fully randomized blocks, although local-control interpolation was used to reduce acquisition-order drift. Bulk viscosity, osmolarity, dissolved oxygen, and water activity were not independently matched across agents, and BSA had a measurable optical background. In addition, the seconds-scale ON/OFF kinetics are integrated fluorescence/protein-state responses and should not be interpreted as radical-pair lifetimes. Replication with independent lysate batches and purified MagLOV2, together with viscosity/osmolarity-matched controls, oxygen control, and structural or time-resolved spectroscopy, will be required to identify the dominant molecular mechanisms.

## 5. Conclusions

Molecular crowding and cosolvent composition strongly regulate MagLOV2 MFE amplitude, recovery kinetics, and pre-field photostability. The distinct responses of Dextran 70, Ficoll PM70, PEG6000, PEG3350, glycerol, ethylene glycol, and BSA show that magnetofluorescence depends on crowder chemistry, polymer size, protein dynamics, and solvent/cofactor environment rather than on a single crowding or viscosity axis. Because MFE amplitude, *τ*_OFF_, and pre-field fluorescence loss are only partially coupled, these readouts provide complementary information about how the molecular environment gates expression of spin-dependent photochemistry. Molecular-environment screening should therefore be considered both a mechanistic probe and an assay-development strategy for identifying and engineering magnetosensitive proteins.

## 6. Author Contributions

Following the CRediT taxonomy [19]:

1. Venkatesh Sridharan: Conceptualization, Data Curation, Formal Analysis, Investigation, Validation, Visualization, Writing-Original Draft
2. Brian L. Ross: Conceptualization, Methodology, Writing-Review and Editing
3. Alessandro Lodesani: Methodology, Resources

## 7. Acknowledgements

We thank Clarice D. Aiello for critically reviewing the manuscript, and Michael Montague for discussions that helped shape the conception of this work. We would also like to thank Morgan Sosa for her assistance with reviewing, formatting, and editing the manuscript.

This work was supported by the Quantum Biology DAO

The [Quantum Biology Institute](http://quantumbiology.org/) is a California non-profit 501(c)(3) focused research organization that performs basic research underpinning the quantum biology field in an open-science fashion.

## Notes

### Competing Interest Statement

The authors have declared no competing interest.

